# Longitudinal network theory approaches identify crucial factors affecting sporulation efficiency in yeast

**DOI:** 10.1101/068270

**Authors:** Camellia Sarkar, Saumya Gupta, Rahul Kumar Verma, Himanshu Sinha, Sarika Jalan

## Abstract

Integrating network theory approaches over longitudinal genome-wide gene expression data is a robust approach to understand the molecular underpinnings of a dynamic biological process. Here, we performed a network-based investigation of longitudinal gene expression changes during sporulation of a yeast strain, SK1. Using global network attributes, viz. clustering coefficient, degree distribution of a node, degree-degree mixing of the connected nodes and disassortativity, we observed dynamic changes in these parameters indicating a highly connected network with inter-module crosstalk. Analysis of local attributes, such as clustering coefficient, hierarchy, betweenness centrality and Granovetter’s weak ties showed that there was an inherent hierarchy under regulatory control that was determined by specific nodes. Biological annotation of these nodes indicated the role of specifically linked pairs of genes in meiosis. These genes act as crucial regulators of sporulation in the highly sporulating SK1 strain. An independent analysis of these network properties in a less efficient sporulating strain helped to understand the heterogeneity of network profiles. We show that comparison of network properties has the potential to identify candidate nodes contributing to the phenotypic diversity of developmental processes in natural populations. Therefore, studying these network parameters as described in this work for dynamic developmental processes, such as sporulation in yeast and eventually in disease progression in humans, can help in identifying candidate factors which are potential regulators of differences between normal and perturbed processes and can be causal targets for intervention.

## INTRODUCTION

Network theory has been used to understand the cellular organization and reprogramming of biological processes (Barabási and Oltvai 2004; Buganim et al. 2012; Gitter et al. 2012). Significant changes in network architecture determined from genome-wide expression studies have identified a few crucial genes such as transcription factors as hubs (Luscombe et al. 2004; Buganim et al. 2012). Transcript abundance of nodes (genes) in a biological network is under genetic control (Brem et al. 2002), and this has led to the understanding of how genetic variation is mechanistically associated with gene expression changes that underlie physiological differences (Ehrenreich et al. 2010; Cubillos et al. 2011). These studies have resulted in the development of exciting approaches aimed at predicting phenotypic consequences of genetic perturbations (Shen et al. 2010). The methods used in these studies perform comparative gene expression analysis such as clustering methods (Eisen et al. 1998), bootstrapping clustering (Kerr and Churchill 2001), four-stage Bayesian model (Wakefield et al. 2003), Gaussian mixture models with a modified Cholesky decomposed covariance structure (McNicholas and Murphy 2010), etc. However, these gene centric methods tend to overlook local patterns where these genes are similar based on only a subset (subspace) of attributes, for example, expression values. This led to an implementation of pattern similarity based bi-clustering approaches to gene expression data that could find bi-clusters among co-regulated genes under the different subset of experimental conditions (Roy et al. 2013).

Sporulation in budding yeast, Saccharomyces cerevisiae, is a linear developmental process initiated under extreme nutrient starvation, involving meiotic cell divisions leading to spore formation (Neiman 2005; 2011). Several genetic, biochemical, and genome-wide transcriptome analyses have elucidated the cascade of transcriptional regulatory processes during sporulation (Fig. 1A). These analyses identified over 1,000 genes that change their expression during meiosis (Chu et al. 1998; Primig et al. 2000). These expression studies categorized the differentially expressed genes into clusters and elucidated their roles in meiosis and sporulation. These studies led to the identification of critical regulatory nodes that are responsible for cells transitioning between different developmental stages during sporulation, viz. *IME1,* initiator of meiosis and *NDT80,* a regulator of meiotic divisions (Chu et al. 1998; Primig et al. 2000). Several sporulation studies have attempted to determine its underlying network structure (Wang et al. 2005; Shen et al. 2010; Ding and Wang 2011). By applying mechanistic models to transcriptional data, for a few crucial genes of sporulation network such as *NDT80*, regulatory mechanisms of this developmental program has been studied (Wang et al. 2005). Most these genome-wide expression studies are done in high sporulation efficiency SK1 strain (90% in 48h, Keeney 2009), while most molecular studies on sporulation are performed on the low sporulating S288c strain (5-10% in 48h; Fig. 1B, Deutschbauer and Davis 2005).

**Fig. 1.**
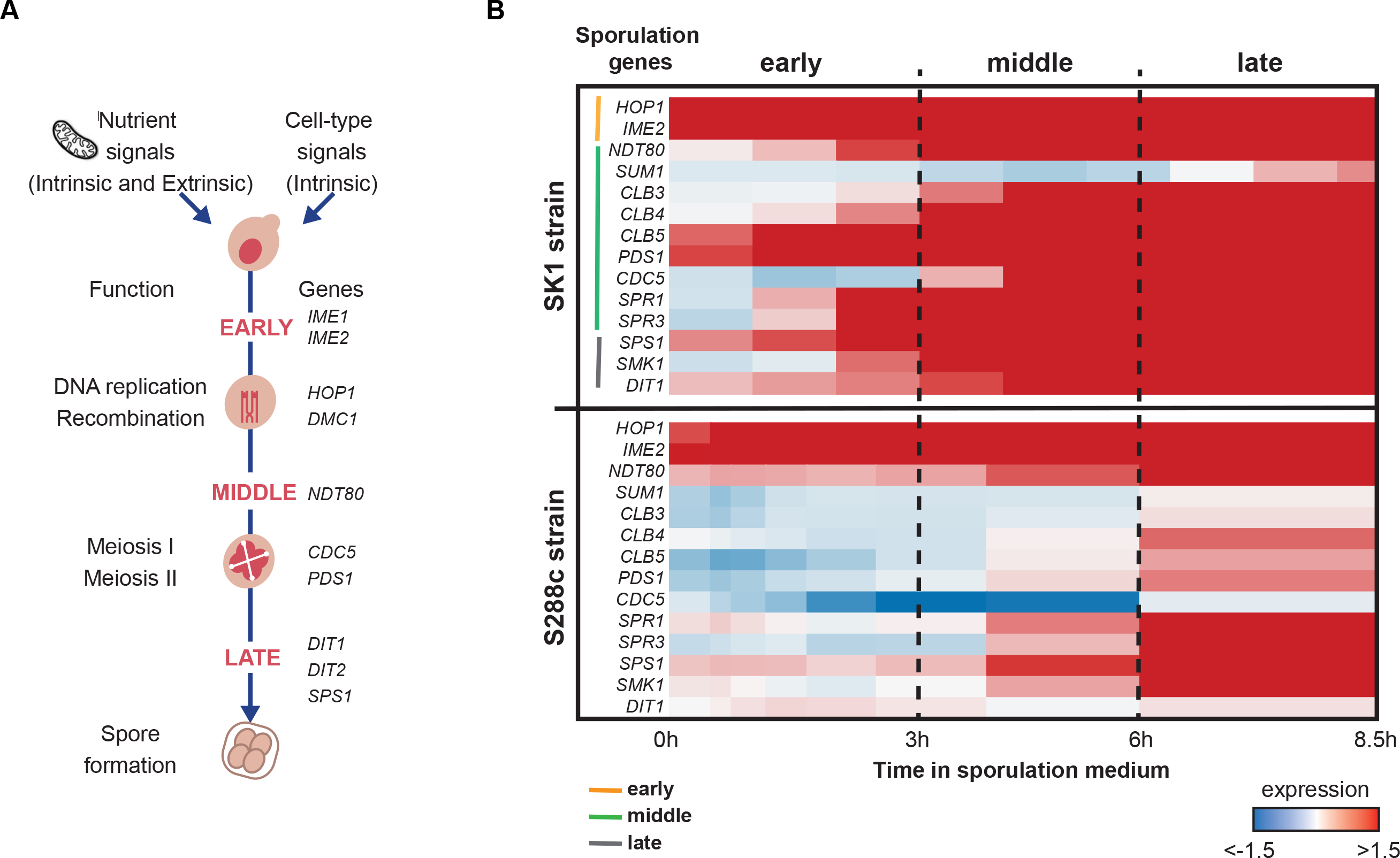
(A) Schematic diagram of the sporulation process. Upon nutrient starvation, the yeast cell exits mitosis (cell division) and initiates meiosis and sporulation. This developmental process of meiosis and sporulation is divided into three phases: early, middle, and late, with each stage having distinct functions and crucial genes activated, adapted from Chu et al. (1998) (B) Heatmap of gene expression profiles of crucial meiosis and sporulation regulators in early, middle and late phases of sporulation. Early, middle, and late phases with their corresponding time points, demarcated by dashed lines, are defined based on the SK1 profile. For SK1, early, middle, and late regulators are indicated by orange, green and grey bars, respectively.

The next step in interpreting gene expression profiles is to go beyond the gene-centric techniques and employ more global approaches for a more comprehensive understanding of how gene expression profiles are specifically related to the regulatory circuitry of the genome (Huang 1999). Network theory provides an efficient framework for capturing structural properties and dynamical behaviour of a range of systems spanning from society (Jalan et al. 2014) to biology (Rai et al. 2014; Shinde et al. 2015). Here we used the network theory approach to investigate how genome-wide transcriptional regulatory networks vary across time and how the determination of various network parameters can help in identifying crucial hubs in this dynamic process. By integrating time-resolved transcriptomics data with the known physical gene interaction network of yeast, we created discrete longitudinal networks of yeast sporulation at multiple time points for the SK1 strain. Using global and local network parameters during sporulation in SK1 strain, we understood its longitudinal networks and identified the nodes that get highly perturbed as this strain enters into the meiotic pathway. An independent longitudinal analysis of these network parameters in the S288c strain that sporulates less efficiently than SK1 was done to understand the heterogeneity of network profiles between these two strains. Identification of crucial nodes and genes helped us to understand how variation in the genetic background can lead to a variable phenotype, sporulation efficiency in this case.

## RESULTS AND DISCUSSION

### Global properties of longitudinal transcriptional regulatory networks of SK1 in sporulation medium

We constructed discrete longitudinal transcriptional regulatory networks of SK1 strain in sporulation medium. This network is composed of 12 sub-networks for each time point during sporulation. Nodes in each sub-network consisted of genes that were differentially expressed with respect to the first-time point (i.e. *t*_0_ = 0h) by at least 2-fold. We first asked how do the different sub-networks of SK1 strain interact with one-another and form functional modules during the sporulation process. For this, we calculated the clustering coefficient for all subnetworks that measures the local cohesiveness between the nodes (Watts and Strogatz 1998). A high value of clustering coefficient of a node depicts high connectivity among its neighbours. We evaluated the average value of clustering coefficient, 〈*C*〉, for each time point. As expected for various biological networks (Albert and Barabási 2002), a high value of 〈*C*〉 was observed for the networks at all time points in SK1 as compared to the corresponding random networks (Fig. 2, Supplementary Table S1) as expected (Watts and Strogatz 1998). To follow these changes, we independently investigated the early (1-4h, as established for this strain previously (Chu et al. 1998; Primig et al. 2000)), middle (4-8h), and late phases of sporulation (later than 8h) in SK1 (Fig. 1B). By comparing 〈*C*〉 the time points, a sharp increase in its value was observed twice for SK1 (Fig. 2B). This increase was at 4h and 9h, signifying modularity in the SK1 network at these time points. Since these early (4h) and middle (9h) time-points are when major meiotic decisions such as those regarding entry into Meiosis I (MI) and Meiosis II (MII), respectively (Chu et al. 1998; Primig et al. 2000) take place, a significant clustering coefficient 〈*C*〉 may play a role in decision making. Furthermore, keeping in view the manner in which we constructed these sporulation networks, a high 〈*C*〉 meant that many of the neighbour target genes of a transcription factor also acted as transcription factors for the other neighbour target genes of that same transcription factor.

**Fig. 2.**
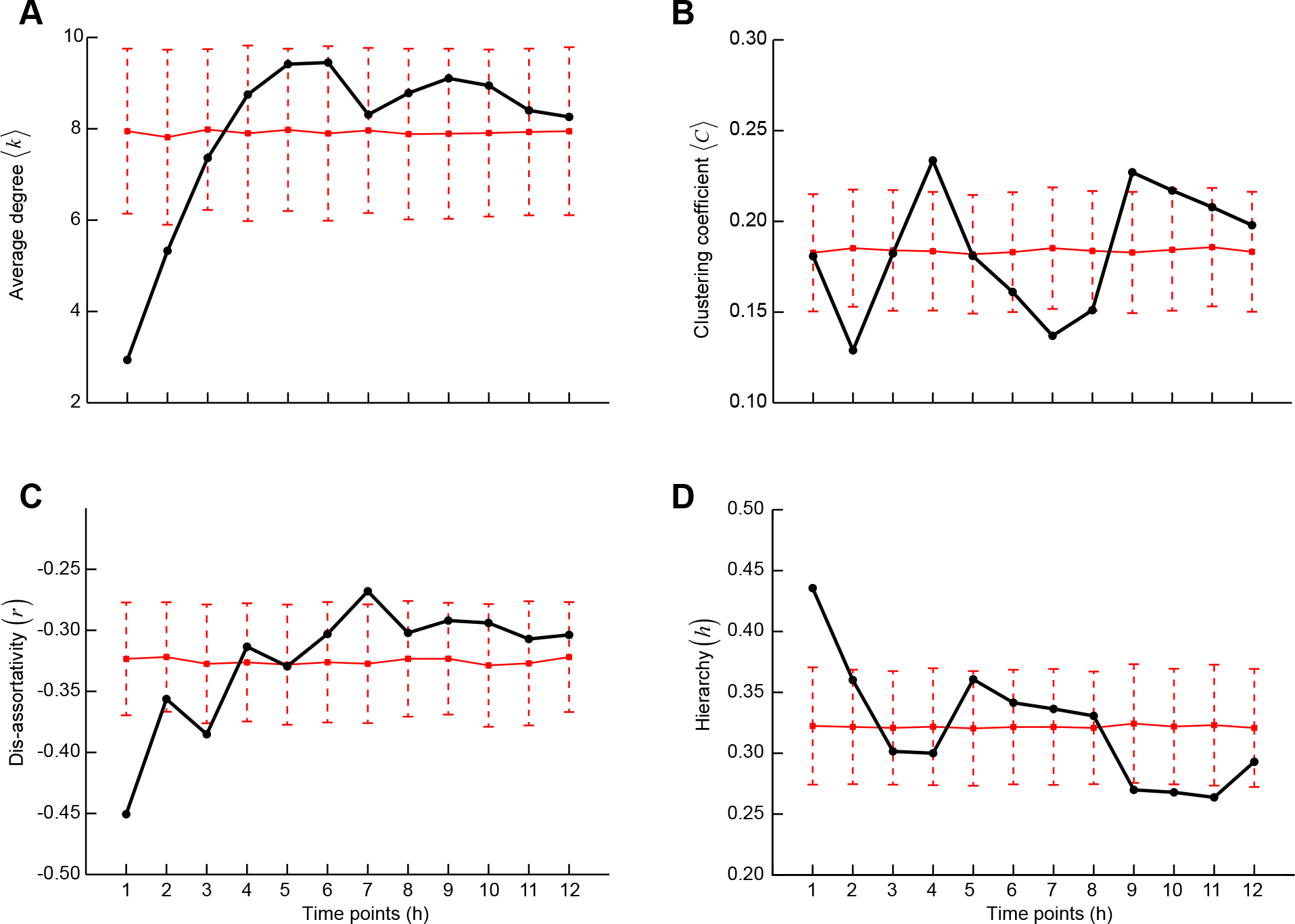
Structural properties of the sporulation networks of SK1 across different time points. (A) Average degree 〈*k*〉; (B) average clustering coefficient 〈*C*〉; (C)Pearson degree-degree correlation coefficient or dis-assortativity (*r*); (D) global reaching centrality or hierarchy (*h*). The solid red line represents the average of 1000 randomised values of each structural property with error bars representing standard deviation.

A closer look at the longitudinal network architecture of sporulation of SK1 revealed that it
showed a heterogeneous degree distribution. Degree distribution 〈*k*〉 is a network parameter defining the number of edges between various nodes. In this biological network, a heterogeneous degree distribution could mean that a few transcription factors dominate the entire network, as is observed in most real-world networks (Albert and Barabási 2002). The sub-networks exhibited a wide range of network sizes across different sporulation time points (Table 1). The larger and denser sub-networks were indicative of extensive regulatory changes in the SK1 strain. We found that there was a drastic increase in the number of genes having significantly high or low expression values in the consecutive time points at the onset of sporulation in the early phase (Table 1), which could be due to cells transitioning from mitotic growth to initiate meiosis. The value of average degree distribution 〈*k*〉 showed a sudden increase in first two points but remained constant across the remaining time points (Fig. 2A). This extensive reprogramming of gene expression early in sporulation as the cells prepare to enter meiotic cell division (Gupta et al. 2016) was revealed as an abrupt increase in the involvement of genes with sporulation progression in the early phase. However, as the sporulation progressed to later phases, the rate of change in network size reduced. Interestingly, despite changes in the early sporulation phase, the ratio of the number of differentially expressed transcription factors (*N*_*TF*_) and target genes remained almost constant across all the time points (Table 1). The proportion of regulatory genes remaining constant throughout sporulation indicated that it might be an intrinsic property of the sporulation process (Supplementary Fig. S1).

**Table 1.**
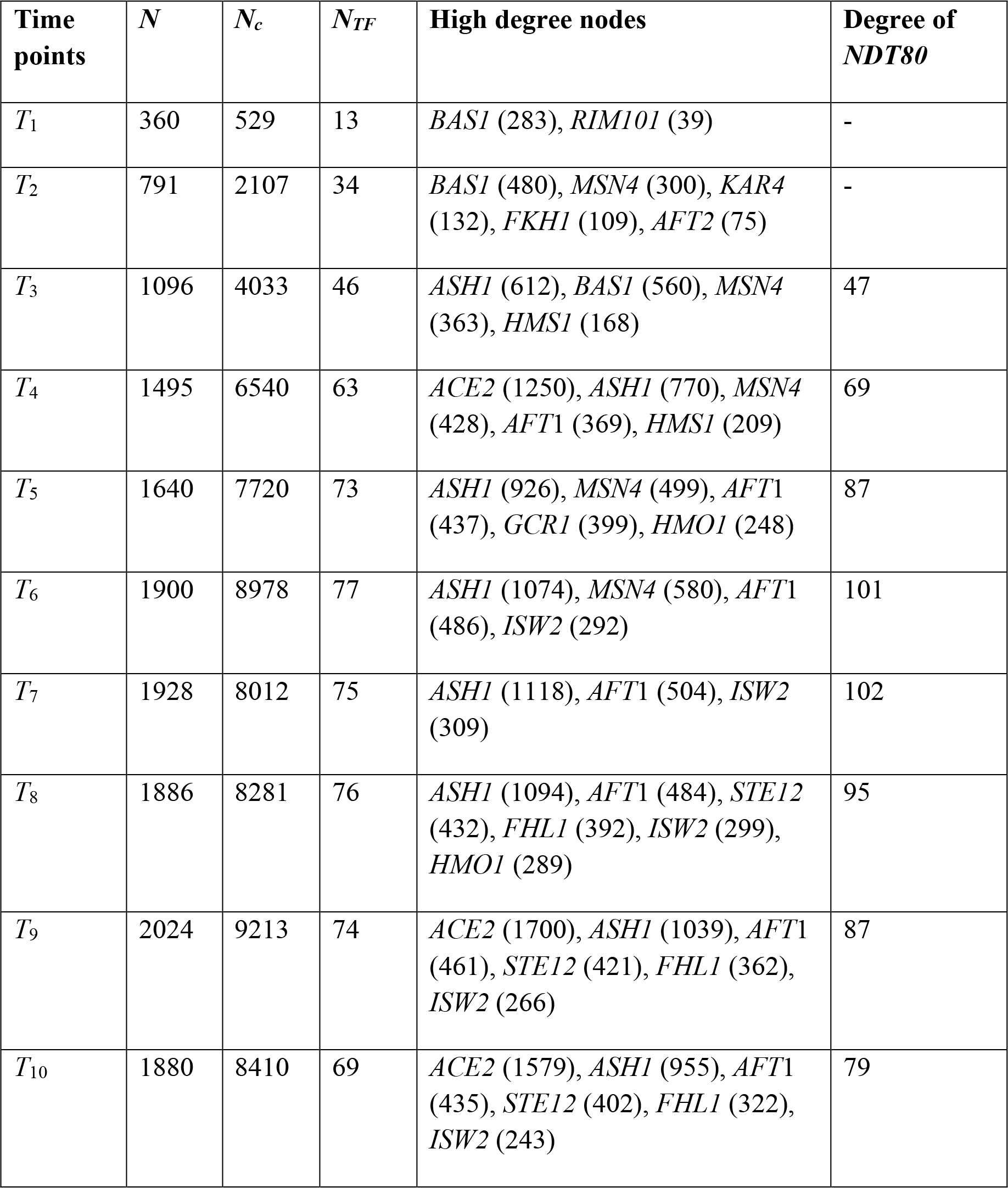
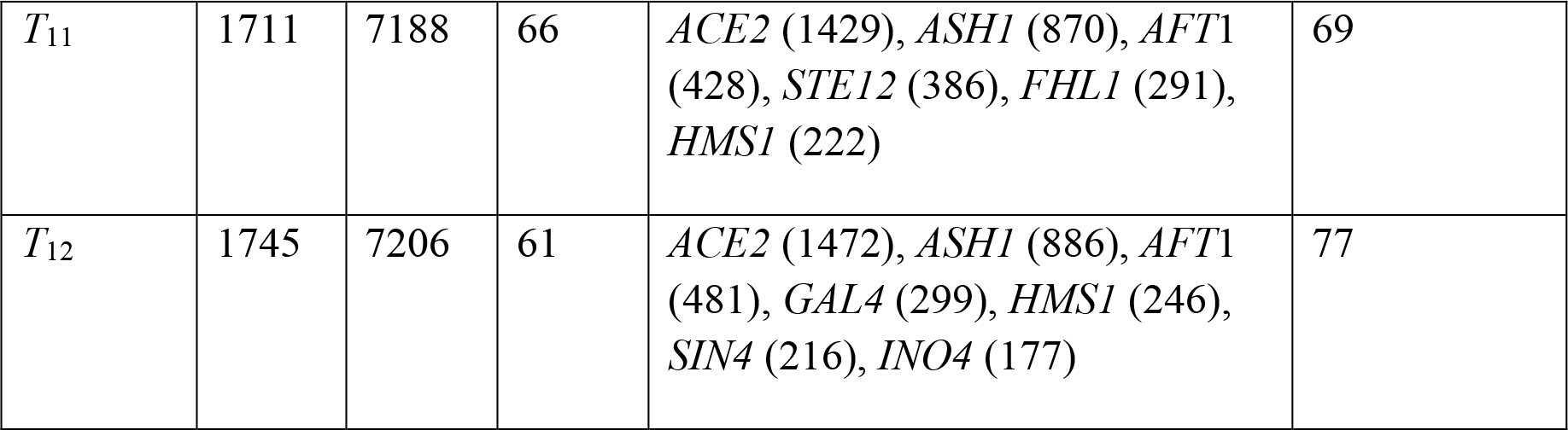
Structural attributes of SK1 networks. *N, N*_*c*_ and *N*_*TF*_ respectively denote the network size, number of connections and number of transcription factors at a particular time point. Catalogue of high degree nodes (degree mentioned in brackets) and degree of *NDT80* gene are given for each time point of SK1 networks.

**Table 2.** Structural attributes of S288c networks. *N, N*_*c*_ and *N*_*TF*_ respectively denote the network size, number of connections and number of transcription factors at a particular time point. Catalogue of high degree nodes (degree mentioned in brackets) and degree of *IME1* gene are given for each time point of S288c networks.

| **Time points** | $N$ | $N_c$ | $N_{TF}$ | **High degree nodes** | **Degree of *IME1*** |
| --- | --- | --- | --- | --- | --- |
| $T_1$ | 434 | 605 | 15 | *BAS1* (356), *HAP4* (92), *TUP1* (48) | 10 |
| $T_2$ | 706 | 1327 | 24 | *BAS1* (523), *STE12* (158), *HAP4* (134), *CUP2* (117), *TUP1* (72) | 14 |
| $T_3$ | 691 | 1280 | 26 | *BAS1* (524), *HAP4* (136), *CUP2* (114), *TUP1* (76), *PUT3* (70) | 13 |
| $T_4$ | 569 | 1025 | 25 | *BAS1* (442), *HAP4* (108), *RLM1* (93) | 13 |
| $T_5$ | 582 | 1049 | 24 | *BAS1* (455), *HAP4* (112), *SWI5* (92) | 15 |
| $T_6$ | 415 | 733 | 23 | *HMO1* (138), *HAP4* (125), *SWI5* (98) | 16 |
| $T_7$ | 409 | 686 | 18 | *HMO1* (131), *HAP4* (124), *SWI5* (97) | 19 |
| $T_8$ | 513 | 1195 | 23 | *MSN4* (261), *HSF1* (246), *HMO1* (129), *SWI5* (95), *INO4* (75) | 21 |

A change in the number of connections modulates the intrinsic properties of a network (Albert and Barabáasi 2002). We investigated the impact of this change on SK1 during the three sporulation phases. Similar to the network size, the number of connections (*N*_*c*_) increased drastically in the early phase. However, the rate of increase in the number of connections was high as compared to the rate of increase in their size, which could be attributed to the appearance of more number of high degree nodes in the second-time point (Table 1). The nodes having high degree refer to genes that regulate a large number of other genes. These highly interacting genes are known to be important in various cellular processes (Rai et al. 2014). In the middle and late phase of sporulation, when processes involved in meiotic divisions occur (Chu et al. 1998), we observed that the number of connections did not show a considerable change since more than 75% of the genes remained same across different time points in this phase (see Supplementary Information).

We next analysed how the interacting patterns impacted the overall structure of the underlying networks. We examined the degree-degree mixing of the connected nodes across the three phases of sporulation in SK1. Disassortativity is a parameter that measures the correlation in the degrees of the nodes in a network and provides an understanding of the dislikelihood in connectivity of the underlying systems (Newman 2002). In gene regulatory networks, highly connected nodes avoid linking directly to each other and instead connect to genes with only a few interactions, thus exhibiting disassortative topology (Yook et al. 2005). This behaviour of the nodes leads to a reduction in crosstalk between different functional modules and increase in the robustness of the networks by localising the effects of deleterious perturbations (Maslov and Sneppen 2002). Pearson (degree-degree) correlation coefficient (*r*) was calculated for the sub-networks at all time points. As expected for gene regulatory networks, sporulation subnetworks in SK1 exhibited disassortativity at all time points (Fig. 2C). A high negative value correlation (*r* = −0.45, Supplementary Table S1) was observed during the early phase of sporulation in SK1 suggesting that the strain was highly resilient to perturbations during early transcriptional events of sporulation (Maslov and Sneppen 2002). After the early phase, in SK1, disassortativity values reach a steady state at middle sporulation phase, suggesting an increase in inter-module crosstalk.

### Local properties of longitudinal regulatory networks of SK1 in sporulation medium

Having analysed the global properties of the sporulation networks, we next studied the impact of the local architecture of SK1 network on the phenotypic profile of the strain. We were interested in investigating how the number of neighbours of nodes denoted by node degree was associated with their neighbour connectivities (interactions between the neighbours of the node of interest) evaluated regarding clustering coefficient. All the networks in SK1 exhibited negative degree-clustering coefficient correlation (Supplementary Fig. S2) as observed in various other real-world networks, indicating the existence of hierarchy in these underlying networks (Barabási and Oltvai 2004). A hierarchical architecture implies that sparsely connected nodes are part of highly clustered areas, with communication between the different highly clustered neighbourhoods maintained by a few hubs. We quantified this hierarchy (*h*), also termed as global reaching centrality in networks (Mones et al. 2012) and found that the networks were more hierarchical at the beginning of sporulation process (Fig. 2D). A high value of hierarchy is associated with modularity in the network. For instance, in case of metabolic networks, hierarchical structure indicates that the sets of genes sharing common neighbour are likely to belong to the same functional class (Ravasz et al. 2002). A low value of *h* indicates more random interactions in the underlying networks. SK1 showed the highest value of hierarchy at time point 1 (*h* = 0.5, around 1h, Supplementary Table S1). A decrease in the hierarchy was observed until the middle phase of sporulation which further diminished in the late phase in SK1, yet again emphasising on the importance of transcriptional regulatory activities in the early phase, on the sporulation process. This inference is in line with the earlier experimental investigations carried out to understand sporulation (Gupta et al. 2015), thus demonstrating the success of this network-based approach in unravelling the underlying genetic basis.

For a network, betweenness centrality is a measure of network resilience (Newman 2001), and it estimates the number of shortest paths (the minimum number of edges traversed between each of the pairs of nodes) that increases if a node is removed from the network (Newman 2006). Usually, nodes with a high degree have high betweenness centrality and are known to bridge different communities in the network. However, in a network, there exist some nodes, which despite having a low degree have relatively high betweenness centrality (Jalan et al. 2014). In the case of gene regulatory networks, such nodes (genes or transcription factors) are involved in less number of regulatory interactions, but these interactions are with different signalling pathways. Thus, these nodes are expected to have particular significance in the underlying networks as their removal can result in a breakdown in the regulatory pathways. Furthermore, in very few cases, a target gene, known to have a low degree may also have relatively higher betweenness centrality than the other target genes if several transcription factors are simultaneously regulating it. For our longitudinal sporulation networks, we identified a few important sporulation-associated genes showing this property (Fig. 3A). These were known regulators of respiratory stress and starvation, namely *STP2* (Merz and Westermann 2009), *PMA1* (Ding et al. 2009) and *RPL2B* (Davey et al. 2012), processes involved in regulation of early phases of sporulation. These results show the importance of this network attribute in identifying nodes with regulatory roles in a dynamic biological process.

**Fig. 3.**
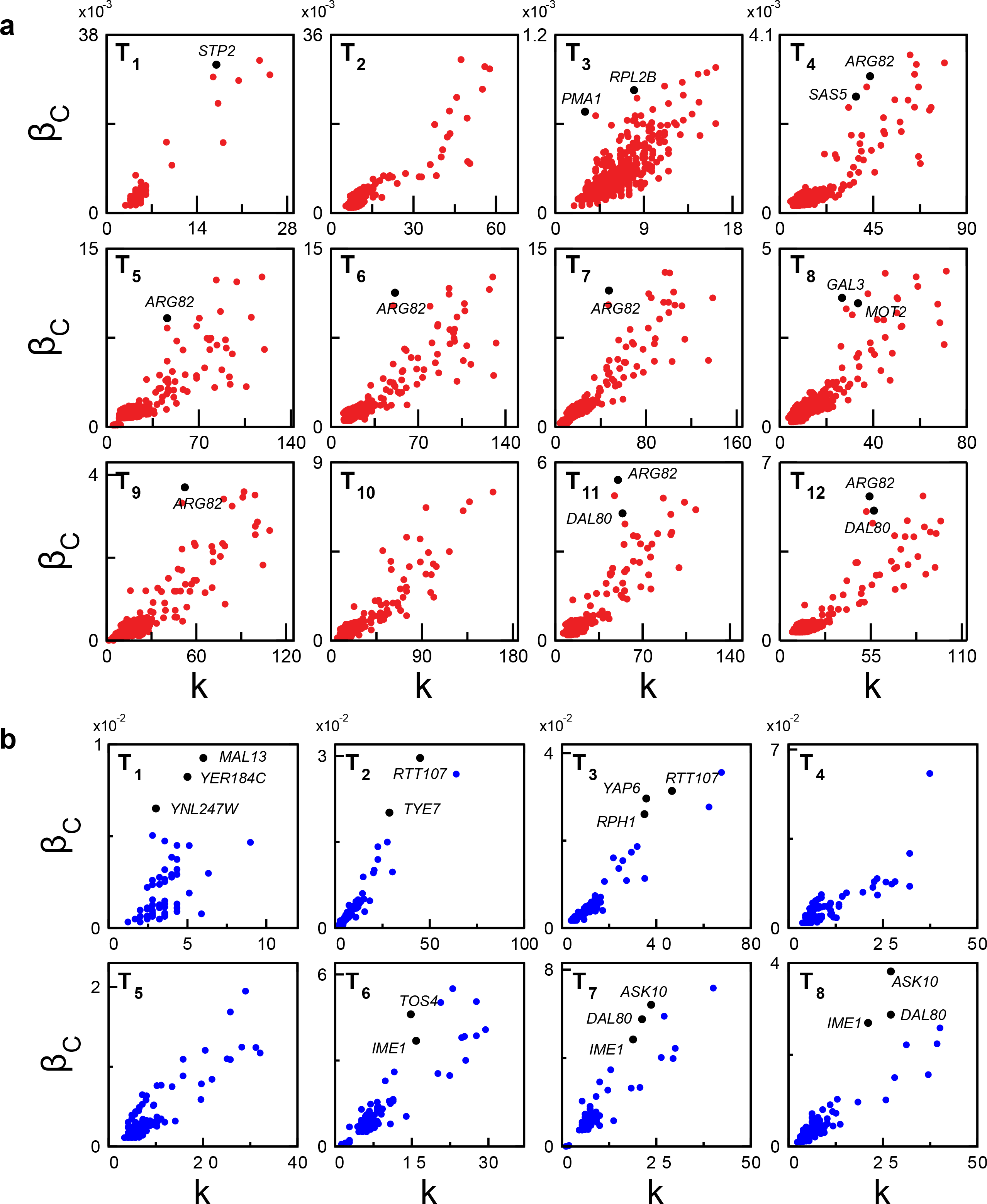
Plots of average degree 〈*k*〉 as a function of betweenness centrality (*β*_*C*_) in (A) SK1 (red) and (B) S288c (blue) networks. Each dot represents a gene and genes with the low degree but higher betweenness centrality in their respective time points are marked as black and named. Supplementary Tables S3 and S4 gives the complete list of genes.

The above analyses helped us to identify influential genes underlying initial transition from mitotic to meiotic cycle which is a significant change in transcriptional profile. We next identified a few interactions that might be instrumental in regulating the sporulation process by considering a significant proposition from sociology, Granovetter’s weak ties hypothesis (Granovetter 1973). This hypothesis states that the degree of overlap of two individuals’ friendship networks varies directly with the strength of their tie to one another. In the networks, the ties having low overlap in their neighbourhoods (i.e. less number of common neighbours) are termed as the weak ties (Onnela et al. 2007). The weak ties that have high link betweenness centrality are the ones known to bridge different communities (Szell and Thurner 2010). Such weak ties revealed through our analysis of different transcriptional regulatory networks are listed in Table 3. Interestingly, we found a repetitive occurrence of the same weak ties in consecutive time points for SK1 indicating their phase-specific importance in yeast sporulation. For instance, *DAL81-ACE2* and *CDC14-ACE2* were repetitive weak ties with high link betweenness centrality in consecutive sub-networks of SK1. To assess the functional importance of these weak ties, we investigated the characteristic properties of the end nodes of these weak ties. Unlike social networks where the end nodes of weak ties are low degree nodes (Sarkar et al. 2016), in the sporulation networks of SK1, the nodes forming weak ties were high degree nodes. An example of this was *BAS1,* a Myb related transcription factor involved in amino acid metabolism and meiosis (Mieczkowski et al. 2006). In addition to *BAS1*, other significant sporulation regulatory genes identified in SK1 were *RIM101,* a pH-responsive regulator of an initiator of meiosis (Su and Mitchell 1993); *IME2,* a serine-threonine kinase activator of *NDT80* and meiosis (Honigberg and Purnapatre 2003); *CDC14*, a protein phosphatase required for meiotic progression (McDonald et al. 2005); *HCM1,* an activator of genes involved in respiration (Rodriguez-Colman et al. 2010) (see Table 3). These results showed that genes showing weak ties and high betweenness centrality are meiosis-associated genes that form essential bridges in SK1. Moreover, *DAL81*, a nitrogen starvation regulator (Marzluf 1997) and *ACE2,* a regulator of G_1_/S transition in the mitotic cell cycle (Spellman et al. 1998) were identified as end nodes of repetitive weak ties in SK1, suggesting their probable regulatory role in the sporulation process that requires further investigation (Table 3).

**Table 3.** Pairs of interacting genes that have low overlap (*O*) and high link betweenness centrality (*β*_L_) in SK1 networks. The interacting gene pairs in bold are mentioned in the main text. Their corresponding indices as given in Supplementary Fig. S4.

| **Time points** | **Node pairs with low $O$ high $\beta_L$** |
| --- | --- |
| $T_1$ | *STP2-BAS1, HCM1-BAS1, BAS1-RIM101* |
| $T_9$ | *ACE2-MGA1, ACE2-SIR2, **DAL81-ACE2**, **CDC14-ACE2*** |
| $T_{10}$ | ***CDC14-ACE2**, ACE2-ARG82, **DAL81-ACE2*** |
| $T_{11}$ | ***DAL81-ACE2**, ACE2-MGA1, **CDC14-ACE2*** |
| $T_{12}$ | *YPL056C-ARG82, **DAL80-ACE2**, RIF1-TPK1* |

**Table 4.** Pairs of interacting genes that have low overlap (*O*) and high link betweenness centrality (*β*_L_) in SK1 networks. The interacting gene pairs in bold are mentioned in the main text. Their corresponding indices as given in Supplementary Fig. S5.

| **Time points** | **Node pairs with low $O$ high $\beta_L$** |
| --- | --- |
| $T_1$ | ***BAS1-RTT107, BAS1-TYE7*** |
| $T_2$ | ***BAS1-RTT107, BAS1-TYE7*** |
| $T_3$ | ***BAS1-RTT107, BAS1-RPH1, YAP6-BAS1*** |
| $T_4$ | ***YAP6-BAS1*** |
| $T_6$ | ***ASK10-HMO1, HMO1-XBP1*** |
| $T_7$ | ***ASK10-HMO1, YAP6-HMO1*** |
| $T_8$ | ***HSF1-IME1, ASK10-HMO1*** |

### Longitudinal transcriptional regulatory networks of S288c in sporulation medium

SK1 is a strain that sporulates efficiently with 90-95% of all its cells entering sporulation within 8-10h in sporulation medium. However, a genetically divergent standard laboratory strain S288c is not an ideal strain to study sporulation as it has low sporulation efficiency (5-10% in 48h; Deutschbauer and Davis 2005). Furthermore, several studies have provided evidence that S288c is not the ideal strain for studying the allelic spectrum in nature, constituting artificial combinations of alleles that have never together been exposed to natural selective pressure (Liti et al. 2009). Whole genome resequencing of 39 yeast isolates was performed to study the genetic variation in genome sequences of diverse strains of yeast, and amongst these strains, S288c diverged strongly by being a phenotypic extreme, across multiple environmental conditions including stresses (Warringer et al. 2011). We were interested in understanding how S288c responds to a stressful environment like sporulation induction, and if by studying global and local network properties, we could identify sporulation-specific differences between S288c and SK1. For this, similar to SK1, we constructed longitudinal transcriptional regulatory networks of S288c strain in sporulation medium that comprised of 8 sub-networks at each time point during sporulation (Gupta et al. 2015). To determine the interactions between these sub-networks and identify the time frame in which S288c forms functional modules, we calculated the average clustering coefficient, 〈*C*〉, for all sub-networks of S288c. While a sharp increase in 〈*C*〉 was observed two times for SK1 coinciding with the early and middle time points in sporulation medium, in S288c, this value was not found to show these characteristic increases (Fig. 4B). A low 〈*C*〉 during middle sporulation in S288c might suggest a lack of interaction between crucial transcription factors and their target genes, leading to low sporulation in this strain. This inability of S288c to form functional modules indicates an inherent property of the strain in responding to stress environments. This inability could be due to a delay in relaying information between functional modules in a cell possibly affecting S288c stress tolerance.

**Fig. 4.**
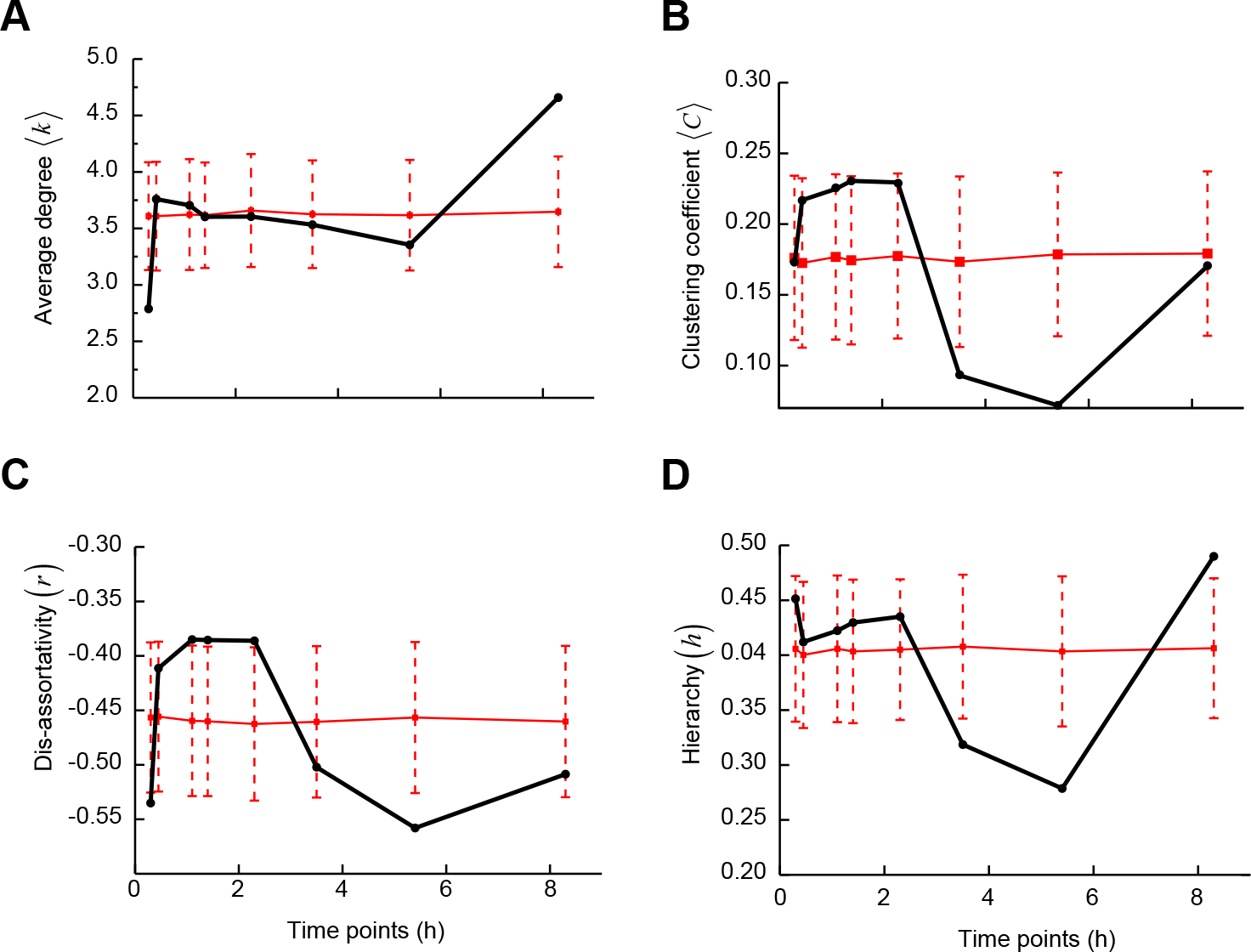
Structural properties of the sporulation networks of S288c across different time points.(A) Average degree 〈*k*〉; (B) average clustering coefficient 〈*C*〉; (C) Pearson degree-degree correlation coefficient or dis-assortativity (*r*); (D) global reaching centrality or hierarchy (*h*). The solid red line represents the average of 1000 randomised values of each structural property with error bars representing standard deviation. Supplementary Table S4 gives the complete list of genes.

When we studied other global network properties of S288c strain, we observed that it exhibited a drastic increase in the number of significantly expressed genes in its early hours in sporulation medium (Table 3) that was similar to SK1. However, the rate of increase in the number of connections (*N*_*c*_) was much higher in the case of SK1 as compared to S288c. For instance, in the early time points in sporulation medium, S288c had a two-fold increase in the number of connections, whereas, SK1 exhibited a four-fold increase (Tables 1 and 2, Fig. 4A). Moreover, the average degree 〈*K*〉 for SK1 was higher in the first few time points compared to S288c with first two time points being significant in SK1 (〈*K*〉 = 2.9 and 5.3) and only first time point in S288c (〈*K*〉 = 3.0, see Figs. 1 and 3, Supplementary Tables S1 and S2).

Since the same repository base network forms the basis for all the interactions for both the strains, a change in the number of connections is possible only if old nodes (genes) disappear or new nodes arise in the networks. Incidentally, as observed in SK1, S288c networks exhibited negative degree-clustering coefficient correlation (Supplementary Fig. S3), indicating the existence of hierarchy in these underlying networks (Barabási and Oltvai 2004). As time progresses, a fall in the number of connections was observed in S288c compared to SK1, with S288c consistently showing lower values for all time points (see Tables 1 and 2). Incidentally, this decrease in the number of connections can be attributed to the disappearance of the high degree node *BAS1,* a Myb-related transcription factor involved in amino acid metabolism and meiosis (Mieczkowski et al. 2006), discussed above as the high-degree end node of weak ties in the SK1 network. Interestingly, in the early time points of S288c, *BAS1* contributed to approximately 50% of the connections (Table 2) even though it is not one of the known regulators of sporulation (Neiman 2011). Its disappearance as time progressed in sporulation medium in S288c was reflected in the number of connections. Furthermore, surprisingly, its involvement in early regulatory processes and disappearance as time progressed, was observed in both the strains. On the one hand, this indicated the specific significance of this gene intrinsic to the early phase of sporulation; on the other hand, it reflected the drastic changes in the regulatory activities as time progresses in sporulation medium. Furthermore, in the later time-points (approximately the time when all the SK1 cells have entered sporulation), the number of connections almost doubled for S288c, and two known stress-responsive regulators, namely *MSN4* and HSF1 (Görner et al. 1998) with a large number of edges appeared in this phase (Table 2). *MSN4*, a known stress response regulator is one of the prime initiators of sporulation by regulating *NDT80.* While it is difficult to speculate whether it is a cause or a consequence - the absence of *MSN4* in the early S288c sub-networks could be one of the reasons behind its poor sporulation ability. This aspect is further exemplified by disassortativity values, which in SK1 was significantly high (close to 0.45) in the early phase of sporulation remaining resilient initially and then a steady increase in module crosstalk. However, in S288c, while the network remained resilient, but module crosstalk fluctuated until very late (Fig. 4C, Supplementary Table S2). This fluctuation implied that the necessary crosstalk between functional modules occurred early in SK1 but was still going on or was random and unstable until later time points for the S288c strain. Furthermore, we observed a significant increase in the hierarchy at around 8h in sporulation medium for S288c, which in comparison was high in early sporulation for SK1 (Fig. 4D).

This late increase in hierarchy again suggested that there was a delay in increased modularity in S288c, which could adversely affect its sporulation process. Crucial regulatory molecular decisions are needed to be taken by a cell in a finite time-window, especially for phenotypes related to developmental processes and stress-related phenotypes. Since metabolic decisions are taking place very rapidly in the early time points of sporulation, if this early signal is relayed later in the S288c, then the finite time-frame is lost. Hence, delayed appearance of crucial regulatory events can be construed as a very critical factor affecting the sporulation process in S288c, and it is possible that the strain remains primarily in the quiescent state resulting in poor sporulation efficiency.

Next, we were interested in identifying influential genes in the S288c longitudinal transcriptional regulatory network in sporulation medium. Interestingly, genes showing the property of low degree and high betweenness centrality were sporulation-associated genes such as *IME1* (Neiman 2011) and *TOS4* (Horak et al. 2002) involved in the initiation of meiosis and DNA replication checkpoint response, respectively. These sporulation-associated genes showed this network property in the early phase in SK1, but in S288c this was only observed at later time points (Supplementary Tables S3, S4). These results again suggested that this late appearance of crucial early sporulation genes as bridges that could transfer information between regulatory modules during early sporulation might be the cause for sporulation not proceeding in S288c. We then examined if identifying instrumental node interactions occurring within 8h in sporulation medium could provide us with candidate molecular pathways that plausibly cause the delay in cross-talk between functional modules of the S288c strain. We observed the repeated occurrence of *BAS1-RTT107, BAS1-TYE7, YAP6-BAS1* and *ASK10-HMO1* in consecutive time-points of S288c sub-networks. We observed apart from *BAS1* (discussed above), genes associated with mitotic functions as end nodes of these weak ties, such as *RTT107* for DNA repair (Leung et al. 2011), *TYE7* for glycolytic gene expression (Sato et al. 2000), *YAP6* for carbohydrate metabolism (Hanlon et al. 2011), *ASK10* for glycerol transport (Beese et al. 2009) and *HMO1* for DNA structure modification (Murugesapillai et al. 2014) (Fig. 3B). While in SK1 meiosis-associated genes formed essential bridges (Fig. 3A), in S288c, genes involved in mitotic functions formed these bridges (Fig. 3B). It implies how differences in weak ties in regulatory networks can help us understand the dramatic differences observed in phenotypes between these strains. It is important to note that since the genome-wide expression data used in these analyses is a bulk expression of a large number of cells, a variation in the number of cells undergoing sporulation, which is the case between SK1 and S288c, can affect the genes identified in our network analyses. Further, it is possible that due to cascading consequences of early dysregulation observed in S288c, many of the following sporulation processes in S288c are dysregulated as well, resulting in changes in network properties compared to SK1. It is not possible to disentangle these two, but our results show that network property analyses were able to identify known differences between SK1 and S288c. Going ahead, it would be interesting to further investigate these genes and their influence on the robustness of the longitudinal network architectures in these strains.

## CONCLUSION

This study presents a novel framework for assessing the molecular underpinnings underlying a phenotype using time-resolved gene expression profile. This framework helped reveal the characteristic signatures of a phenotype and identified candidate genes contributing to and affecting the phenotype. Inferences drawn based on the comparative investigations of structural attributes of the longitudinal sporulation networks of the two strains revealed that late appearance of early regulators and delayed crosstalk between functional modules might be critical for progression of sporulation process in SK1 and the plausible reasons behind the low sporulation efficiency of S288c. This speculation is especially interesting since most causative genetic variants known to contribute to sporulation efficiency variation have been observed in genes either showing early role in sporulation or affecting genes with an early regulatory role in sporulation (Deutschbauer and Davis 2005; Ben-Ari et al. 2006; Gupta et al. 2014; 2016).

Application of genome-wide strategies to elucidate the molecular networks in multiple genetic backgrounds provides us with the opportunity to understand the impact of natural variation. Studying these network properties for variation in causal genes would further help in understanding specific molecular effects in the different temporal phases of the phenotype. The strategies adopted in this work can be extended to assess the impact of molecular perturbations in the already known core interaction network of an organism (Carter et al. 2013; Gasch et al. 2016). Moreover, application of such network analyses on gene expression datasets for disease progression in complex diseases such as cancer and metabolic disorders can help identify specific nodes perturbing the underlying molecular pathways that can be the focus of personalised medicine and drug target discovery.

## METHODS

### Network construction

For constructing the transcriptional regulatory sporulation network, the known static regulatory interactions were overlaid on the time-resolved transcriptomics data of the two strains. The overlaying of the regulatory network and temporal transcriptome data created the longitudinally integrated sporulation networks. The static network known for yeast contains all the known regulatory interactions between all the yeast transcription factors (TF) and their target genes (TG). These interactions were obtained from YEASTRACT database (Teixeira et al. 2013), a curated repository of regulatory associations in S. cerevisiae, based on more than 1,200 literature references.

Gene expression data were obtained from previously published studies - for SK1 (Lardenois et al. 2011), and S288c (Gupta et al. 2015). These datasets contained gene expression of 6,926 genes across 13 different time points in linear scale (0h to 12h with 1h intervals termed as T0 to T12, respectively) in SK1 and 9 different time points in logarithmic scale (0h, 30m, 45m, 1h10m, 1h40m, 2h30m, 3h50m, 5h40m, 8h30m termed as T0 to T8, respectively) in S288c. Gene expression analysis was performed as described previously (Gupta et al. 2015). In brief, all time points were normalized together using vsn (Huber et al. 2002) and the log2 transformed expression values obtained after normalisation were smoothed using locfit. Fold differences in expression values were calculated for all the time-points relative to t = 0h (t0), as follows:

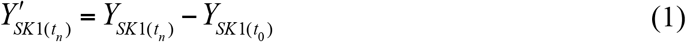

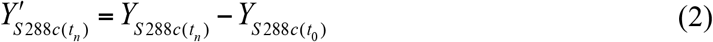

such that *Y* is the expression value of a transcript for a strain (SK1 or S288c) at a specific time point *n* and *Y’* is the transformed expression value. One of the reasons behind considering fold-change values is to mask the effect of any possible transcriptional noise arising in the system.

At each time point, differentially expressed genes were identified by setting the threshold value on *log*_*2*_ fold differences as 1.0. Hence, genes that were considered over-expressed or repressed showed at least a 2-fold difference with respect to the first time point *t*_0_ (i.e. t = 0h) (Gupta et al. 2015). This step again reduces the possibility of picking up any gene that contributes to transcriptional noise.

The longitudinal sporulation network was constructed by overlaying the experimentally determined yeast sporulation-specific gene expression values on the yeast static network. For each time point of each strain, only those TF-TG pairs were considered that both showed either overexpression or repression. These pairs were included in the subnetwork for that specific time point and thus, subnetworks for each time point were constructed for each strain. For comparison of the gene names obtained from YEASTRACT and the sporulation gene expression data, aliases were obtained from Saccharomyces Genome Database (Cherry et al. 2012). This step again refines the interaction data.

### Data availability

The codes, supplementary tables and figures, adjacency matrices of the networks constructed using longitudinal sporulation data drawn from SK1 and S288c strains, the corresponding gene indices and transcription factors are freely available online at figshare https://doi.org/10.6084/m9.figshare.3457508.v4.

### Structural parameters

Several statistical measures are proposed to understand specific features of the network (Albert and Barabási 2002; Boccaletti et al. 2006). The number of connections possessed by a node is termed as its degree. The spread in the degrees is characterised by a distribution function *P* (*k*), which gives the probability that a randomly selected node has precisely *k* edges. The degree distribution of a random graph is a Poisson distribution with a peak at *P*(〈*k*〉). However, in most large networks such as the World Wide Web, the Internet or the metabolic networks, the degree distribution significantly deviates from a Poisson distribution but has a power-law tail *P*(*k*)~ *k*^−γ^. The inherent tendency of social networks to form clusters representing circles of friends or acquaintances in which every member knows every other member is quantified by the clustering coefficient. We categorise the nodes as high and low degree nodes by arranging all the nodes in a network in descending order of degrees and keep assigning the nodes as high degree nodes until the next lower degree node differs by nearly 1.5-fold from the former in terms of the degree. The clustering coefficient of a node *i*denoted as *C*_*i*_, is defined as the ratio of the number of links existing between the neighbours of the node to the possible number of links that could exist between the neighbours of that node (Newman et al. 2001) and is given by

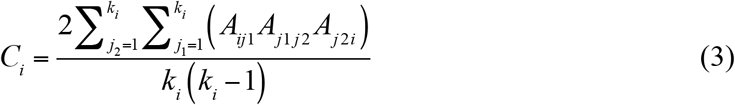

where *i* is the node of interest and *j*_1_, and *j*_2_ are any two neighbours of the node *i* and *k*_*i*_ is the degree of the node *i*. The average clustering coefficient of a network corresponding to a particular condition 〈*C*〉 can be written as

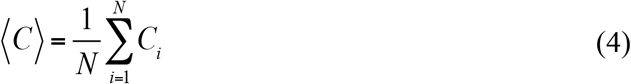

We define the betweenness centrality of node *i*, as the fraction of shortest paths between node pairs that pass through the said node of interest (Newman 2006).

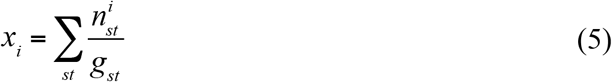

where *n*_*i*_ is the number of geodesic paths from *s* to *t* that passes through *i* and *g*_*st*_ is the total number of geodesic paths from *s* to *t*. All the nodes were plotted, and the top 5% of the nodes (genes) with high betweenness centrality but low degree were identified.

We quantify the degree-degree correlations of a network by considering the Pearson (degree degree) correlation coefficient, given by Newman (2002)

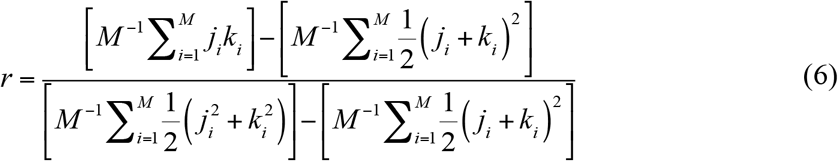

where *j*_*i*_, *k*_*i*_ are the degrees of nodes at both the ends of the *i*^*th*^ connection and *M* represents the total connections in the network.

Link betweenness centrality is defined for an undirected link as

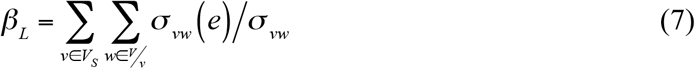

where *σ*_*vw*_ (*e*) is the number of shortest paths between *v* and *w* that contain *e*, and *σ*_*vw*_ is the total number of shortest paths between *v* and *w* (Onnela et al. 2007).

The overlap of the neighbourhood of two connected nodes *i* and *j* is defined as (Onnela et al. 2007)

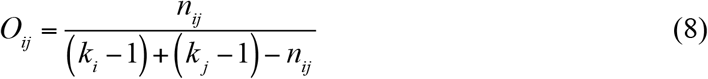

where *n*_*ij*_ is the number of neighbours common to both nodes *i* and *j*. Here *k*_*i*_ and *k*_*j*_ represent the degree of the *i*^*th*^ and *j*^*th*^ nodes.

The hierarchy can be defined as the heterogeneous distribution of local reaching centrality of
nodes in the network. The local reaching centrality, (*C*_*R*_), of a node *i* is defined as (Mones et al. 2012)

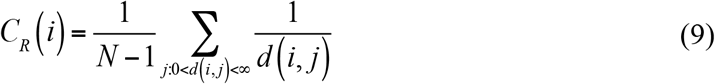

where *d*(*i,j*) is the length of the shortest path between any pair of nodes *i* and *j*. The measure of hierarchy (*h*), termed as global reaching centrality is given by

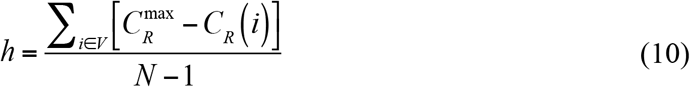

## ACKNOWLEDGEMENTS

SJ acknowledges Department of Science and Technology (DST), Govt. of India grant EMR/2014/000368 and Council of Scientific and Industrial Research (CSIR), Govt. of India grant 25(0205)/12/EMR-II. SG acknowledges support by Tata Institute of Fundamental Research during the initial phase. HS acknowledges support by Tata Institute of Fundamental Research during the initial phase and Indian Institute of Technology Madras.

## AUTHOR CONTRIBUTIONS STATEMENT

SJ conceived the idea. SJ and HS designed and supervised the project. CS, RKV constructed the networks with the help of SG and analysed the structural properties. SG, RKV, CS, HS analysed the functional properties. CS, SG, HS wrote the initial draft, and all the authors contributed to the manuscript.

## SUPPLEMENTARY INFORMATION

Supplementary information accompanies this paper.

## COMPETING FINANCIAL INTERESTS STATEMENT

The authors declare no competing financial interests.

